# Comparative performance analysis of mulberry silkworm (*Bombyx mori* L.) races for integration of superior yield and maximize profit in West Bengal

**DOI:** 10.1101/2021.03.22.436391

**Authors:** Megha Kaviraj, Upendra Kumar, SN Chatterjee, SL Swain, M Singh, Kasinath Nandi

**Affiliations:** ICAR-National Rice Research Institute, Cuttack, Odisha -753006, India; The University of Burdwan, Burdwan, West Bengal-713104, India; Fakir Mohan University, Balashore, Odisha-756020, India; Triveni Devi Bhalotia College, Raniganj, West Bengal-713347, India; Asansol Girl’s College, Burdwan, West Bengal-713304, India

**Keywords:** *Bombyx mori*, Bivoltine, Multivoltine, F1 variety, Economic traits, Heatmap

## Abstract

India has a prosperous and glorious history in silk production and its silk trade dates back to 15^th^ century. Sericulture is practiced in many regions of India, where West Bengal ranks third in mulberry silk production. About 2000 villages are engaged in mulberry cultivation with plantation area of 37,883 acres. But nowadays, farmers have lack their interest in this sector due to low cost benefit ratio, high investment cost in terms of silk worm rearing, land uses and lack of proper knowledge on different races of *Bombyx mori*. Therefore, the present study has been aiming to find out the comparative performance of bivoltine, multivoltine, and F1 variety of *B. mori* by analysed the cocoon, post-cocoon characters, and silk related traits for identifying the best superior races of *B. mori*. The results showed that feeding habits of silkworm larvae of three races had significant effect on cocoon parameters. The average weight of ten 5th instar larvae of bivoltine was higher in comparison with multivoltine and F1 variety. Similar trends were obtained in the average length of silk thread from a single cocoon in bivoltine races, higher by 30% and 77.14% from F1 and multivoltine, respectively (P<0.05). We also inquired about capital investment and profit-making of three varieties. Net profit for bivoltine was significantly increased by 15.65% and 10.21% than multivoltine and F1 varieties, respectively.The heatmap analysis revealed that the bivoltine races of *B. mori* were separately clustered based on the positive correlation of measured variables mostly length of silk thread, renditta, dry cocoon weight, the average weight of larvae, present market price, leaves for rearing, net profit and selling price of cocoon per acre farmland; whereas, the multivoltine and F1 variety clustered in a separate group. From our findings, it is clear that the culture of bivoltine is preferable for the farmers who don’t have enough landed property and also profitable in terms of cocoon, post cocoon and silk traits. Therefore, it is recommended that the farming of bivoltine silkworm is more gainfulcompared to F1 and multivoltine variety, and capable of more income generation than other traditional agricultural crops.

**Fig.**
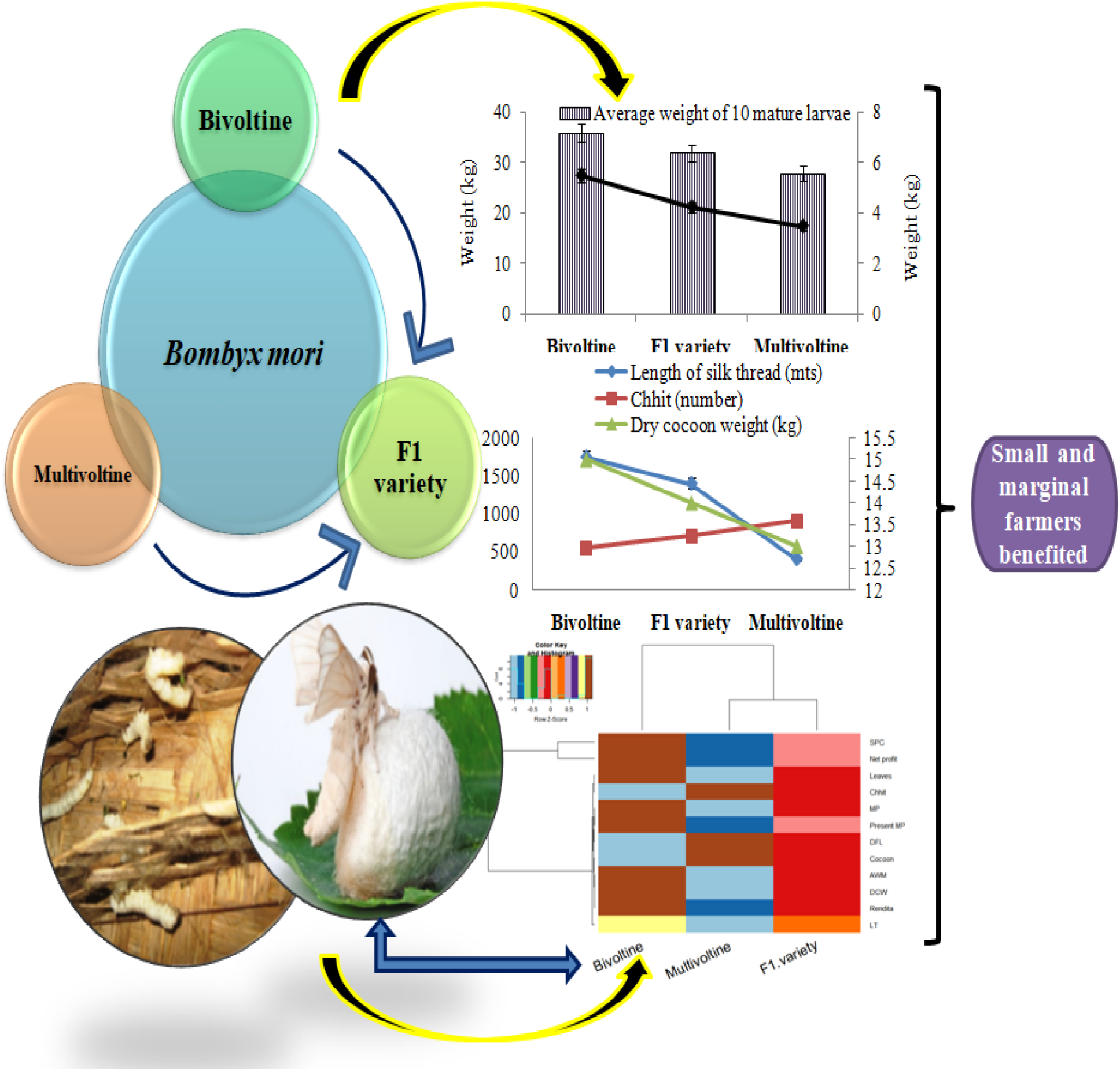
Graphical abstract represents the comparative performance of bivoltine, multivoltine and F1 races of *B. mori* on the aspect of biological and economic traits in tropical climate condition.

## 1. Introduction

India is the 2^nd^ largest raw silk manufacturer country and produces five commercially significant silk varieties namely, Mulberry, Tropical Tasar, Temperate Tasar, Eri and Muga (CSB, 2018). It is one of the vital sub-sectors of agriculture and plays a crucial role in farm economy through providing employment round the year and fetches higher income mainly to the rural farm families (Roopa and Murthy, 2015; Masrat and Tripathi, 2017). Among all silk varieties, mulberry silk occupies 75% of the total silk production. Sericulture is practiced in many states of India, West Bengal ranks third in mulberry silk production and is highly concentrated in three traditional sericulture districts namely, Malda, Murshidabad and Birbhum, which contributing 90% of the total state’s silk production. Currently,West Bengal accounts for 14.5% of the total country’s cocoon production (CSB, 2018) and about 2000 villages are directly involved in mulberry cultivation with plantation area of 37,883 acres (Anonymous, 2016).

Rearing of silkworm on large scales with great care in both natural and control conditions scientifically for cocoon production is used as the raw material for silk production (Kamili et al., 2000). Silk is popularly known as “Queen of Textiles” in all over the world for the nature of its unparalleled grandeur, sheen, inherent affinity for dyes, high absorbance, light weight, soft touch and high durability. Mulberry silkworm *Bombyx mori* (Lepidoptera: Bombycidae) spins the precious silk fibre, which making it one of the most beneficial insects to mankind, and is becoming an attractive multifunctional material for both textile and nontextile uses (Tsukada et al., 2005). Approximately all-commercial silk is made from cocoons spun by silkworms of the genus *Bombyx* (Lee, 1999).

Mulberry (*Morus* L., Family Moraceae), is an essential foliage crop in sericulture as it is the only natural feed of the silkworm *Bombyx mori*. Previously, it is believed to have originated at the Himalayans foothills (Koidzumi, 1917). Earlier, Linneaus (1753) classified the genus *Morus* into five species on the basis of morphological characteristics: *Morus alba, M. nigra, M. rubra, M. tartarica* and *M. indica* L. The quality of the mulberry leaves has a direct impact on the normal growth and development of the larvae and also the quality of the cocoon (Legay, 1958; Adolkar et al., 2007; Gangwar, 2010). Different races of silkworms are characterized by undergoing one, two or several uninterrupted generations during the summer. Such races are called uni-, bi- or multivoltine.

Though, India is the second largest producer of raw silk in the world next to China with an annual production of 31,931 MT (CSB, 2018) but the raw silk yarn is of low standard due to multivoltine in nature. Besides, other reason behind low standard of silk is the tropical climatic conditions of the country with marginal sub-tropical and temperate sericultural areas. In tropical areas of the country, multi x bi hybrids are reared and produced silk is not superior in quality and as such is not popular at International market. To overcome this drawback, compatible bivoltine breeds/hybrids for rearing under tropical conditions were developed (Lakshmi et al., 2011) and selected for rearing in field conditions. The productivity of sericulture mainly relies on high breeding stock of the silkworm. The hybridization is a technique used to enhance the yield of silkworm and cocoon production (Brahma et al., 2015). By crossing genetically distinct population understanding the genetic mechanism of the silkworm, high yielding and disease tolerant races with distinct quantitative and qualitative traits can be achieved. The success in silkworm hybridization primarily depends on the selection of initial breeding materials followed by their effective utilization in different combinations to create genetic variability for selection (Mano et al., 1992). In India, it is estimated that nearly 80% of the silk is produced by multivoltine × bivoltine hybrids where multivoltine races are used as a female parent for commercial exploitation. The main reason attributed to this is that the contribution of bivoltine by virtue of its maternal inheritance may result in regular crop loss (Doddaswamy et al., 2009). Suitable silkworm hybrids play a vital role in increasing the productivity and quality of silk which are important for sustainable sericulture industry (Kumar et al., 2011). As per available literature, manifestation of heterosis in silkworm has been established by many breeders (Talebi and Subramanya, 2009; Kumar et al., 2012). A variety of hypotheses have been advocated as to the inheritance of voltinism.However, information on three races of *B. mori* in different voltinism pattern is scanty.

Hence, in the present study, an attempt has been made to analyze the comparative performance of bivoltine, multivoltine and F1 variety of*B. mori* with significant replications and also, analyzedthe cocoon post-cocoon characters and silk related traits for further commercial application of the best superior races of *B. mori*.The purpose of this study was to provide information about the superiority based on the aspect of biological and economical perspective of three races of mulberry silk worm for researcher as well as for farmer’s community for sustainable income generation.

## 2. Methodology

### 2.1 Experimental site

The present study was conducted at Bishnupur sericulture complex, Bankura, West Bengal, which is located between 23°03□25□N and 87°18□15□E. The altitude is 59 m above the sea level. Bishnupur’s climate is classified as tropical (Fig. 1). The average temperature was 26.6°C and about 1157 mm of precipitation falls annually. The lowest (2 mm) and greatest (255 mm) precipitation was occurred in the month of July and December, respectively.

**Fig. 1.**
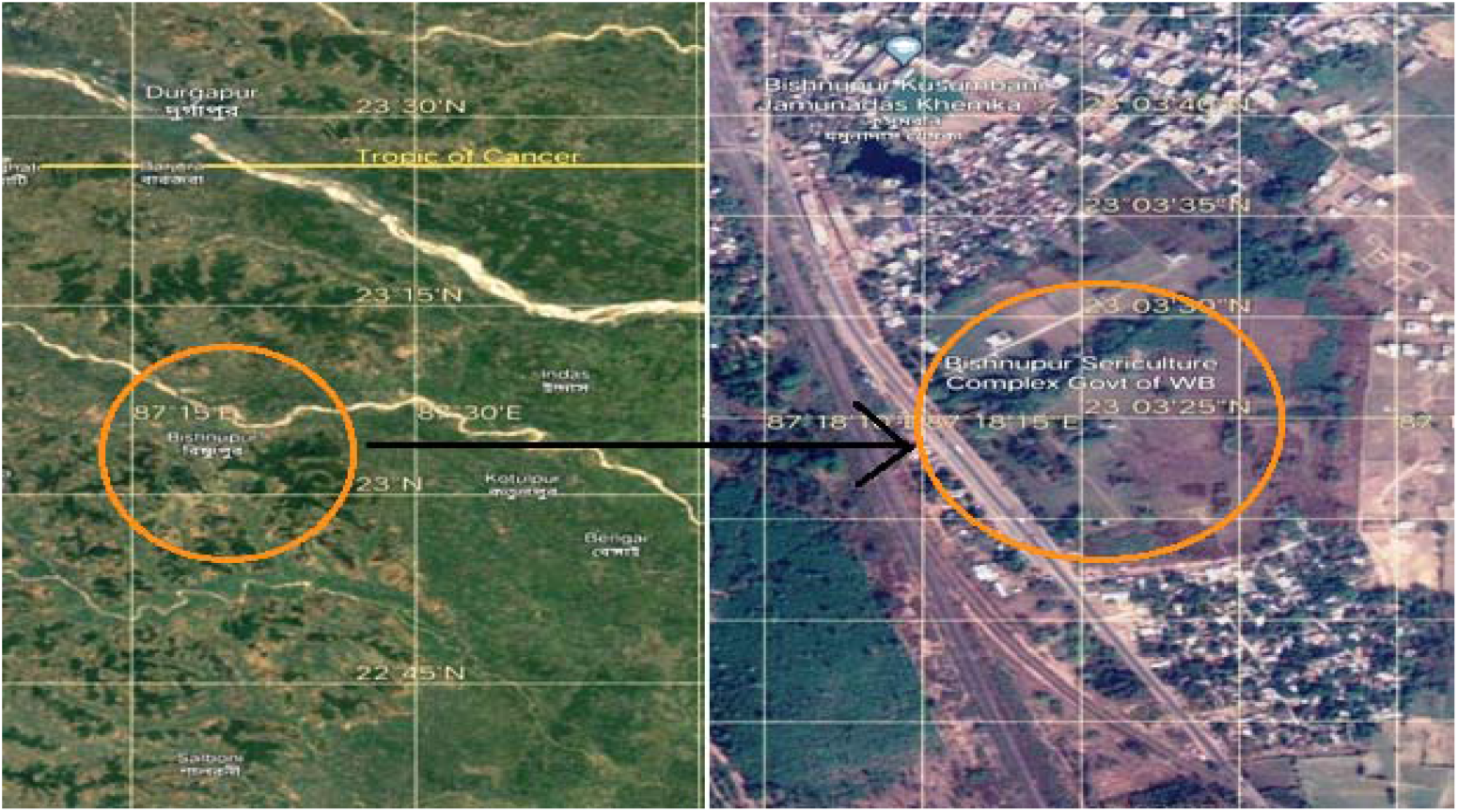
Geographical location and coordinates of experimental site in Bishnupur sericulture complex, Government of West Bengal at Bishnupur, West Bengal, India.

### 2.2 Experimental material and set up

Disease-free egg layings (dfls) of the bivoltine (AP8), multivoltine (Nistari) and F1 variety (a hybridized variety by crossing the multivoltine female moth with bivoltine male moth and the eggs laid after mating are termed as F1 variety) of silkworm strains of *Bombyx mori* L. were used in this investigation. Different races of silkworms are characterized by undergoing one, two or several uninterrupted generations during the summer; such races are called uni-, bi- or multivoltine.Rearing of all the silkworms was done by adopted the procedures of Jolly (1987) and Ullal (1987). Silkworms were reared under standard conditions of 26 ± 2°C, 70 ± 5% relative humidity and 12:12 (L:D) photoperiod (Raina, 2000) (Table 1). Rearing was done in trays measuring 90 x 60cm, placed on rearing racks, 150 x 75 x 200cm that could hold 24 trays each. The silkworm larvae were reared in these trays from their first instar to the fourth. At the onset of the fifth instar, 100 worms of each strain were randomly selected and monitored individually (Fig 2b, 2c). Temperature and humidity in the room ranged between 24 - 27°C and 84% - 86% respectively for young age rearing and 23 - 24°C and 65% - 70%, respectively for late age rearing. The rearing room was white-washed and fumigated with sulphur dioxide gas. Rearing trays, stands, incubator and all other tools were disinfected with Formaldehyde (2%) solution. Data was recorded replication-wise for all three variety viz., fecundity, hatching percentage, larval weight, larval survival percentage, cocoon yield per 10,000 larvae (by weight and by number), good cocoon percentage, pupation percentage, single cocoon weight, single shell weight, average filament length adapted from standard protocol.

**Fig. 2.**
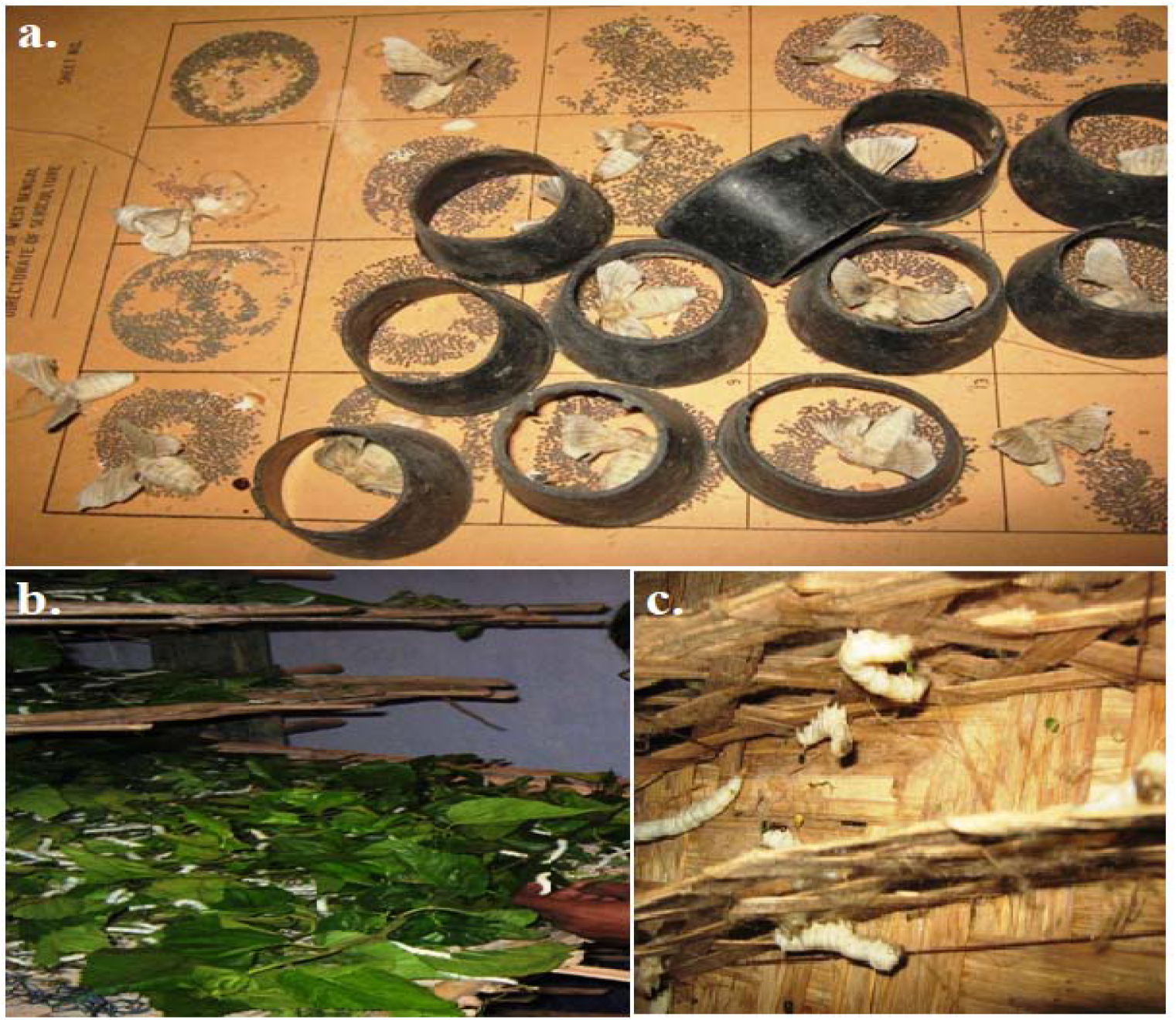
**a.** Photograph of cellule on the card within which the female moths are laying eggs. b.Mature fifth instar larvae of M12W multivoltine variety feeds on fresh mulberry leaves. c. Mature larvae of multivoltine variety just starting cocoon formation on chandraki.

**Table 1.** Environmental condition required for larval growth of silk worm.

| Larval stage | Temperature | Humidity |
| --- | --- | --- |
| 1<sup>st</sup> inster larva | 26-28°C | 85% |
| 2<sup>nd</sup> inster larva | 26-28°C | 85% |
| 3<sup>rd</sup> inster larva | 24-26°C | 80% |
| 4<sup>th</sup> inster larva | 24-25°C | 75% |
| 5<sup>th</sup> inster larva | 23-24°C | 70% |

### 2.3 Food plant and consumption

Mulberry silk worm, being monophagus insect, feeds only on mulberry (*Morus indica)* leaves. At present mulberry is cultivated in about 2.90 lakh hectares in India. Generally, the favorable growth temperature of mulberry plant is ranging from 24°C - 35°C and prefers slightly the acidic soil ranging from pH 6.2 - 8.2. The favorable plantation season is early monsoon. However, if proper irrigation facilities are available, mulberry can be planted at any time. Mulberry grows in a wide range of soil from clayey loam to loamy soil. Mulberry is a deep rooted perennial plant, therefore, the soil having good water retention capacity. The larvae were fed with mulberry leaves (Fig. 2b). Proper and adequate food provided to the larvae because it greatly influence the quality of cocoon. Feeding must satisfy both the appetite and nutritional requirements of the larvae. Normally 4 - 6 feedings were provided per day.

### 2.4 Weight of 10 mature larvae

Ten mature larvae were picked randomly from each replicate from fourth to sixth day of final instar and weighed on digital balance and represented as a unit of ‘g’. Maximum larval weight was recorded in each hybrid combination (Jolly, 1987).

### 2.5 Quantity of leaves required for larvae rearing

Amount of leaves required for the rearing of 100 disease free larvae (DFL) of different three races of *B. mori* and the weight of the leaves expressed in kg.The larvae to be reared should be free from any sort of disease or in other words the eggs should be disease free laying (DFL).

### 2.6 Cocoon characters

Following observations were made for different parameters at cocoon stage viz., effective rate of rearing (ERR) by green and dry cocoon weight and by number (Radjabi, 2011).

#### 2.6.1 Chhit /green cocoon weight

The weight of freshly produced cocoon within which the larvae or pupae are in living condition called chhit. In sericulture industry the term chhit implies the number of green cocoons weigh in one kilogram weight.

#### 2.6.2 Dry coccon weight

The weight of cocoon which was completely dried either in sun or electric oven.The dry cocoon weight of 40 Kg green cocoon from different varieties of mulberry silk worm was recorded.

### 2.7 Post cocoon characters

After harvesting, cocoon samples were stifled in hot air oven at 90-600°C for six hours forreeling purpose. Randomly selected cocoons were reeled for post cocoon parameter and data of different parameters viz., total filament length, non-breakable filament length, denier and renditta was recorded (Ullal, 1987).

#### 2.7.1 Total filament length

It represents the total length of silk filament reeled from a single cocoon in meters randomly selected. Ten cocoons from each replicate were reeled and average filament length was calculated as:

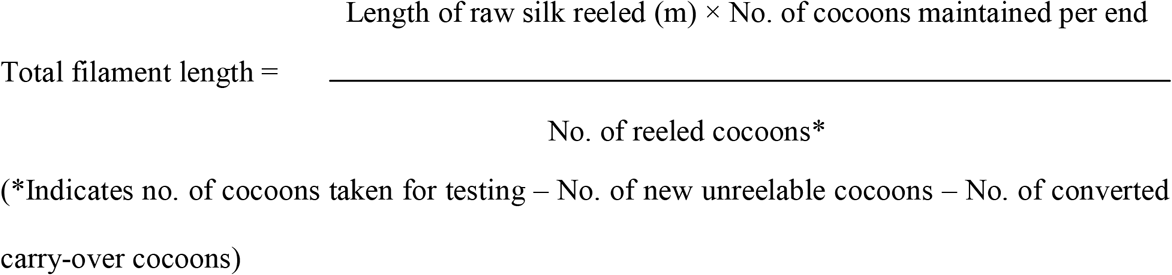

(*Indicates no. of cocoons taken for testing – No. of new unreelable cocoons – No. of converted carry-over cocoons)

#### 2.7.2 Renditta

Renditta is the amount of cocoon required to extract one kg. of silk thread. Say for example if 10 kg. of cocoon is required to extract 1 kg. of silk thread, its renditta will be 10. Here, we estimated the average amount of silk thread produced from 40 kg. of different varieties of cocoons.

### 2.8 Number of DFL reared and cocoon produced in per acre farm land

The mating is done in captivity located at Bishnupur sericulture farm for both bivoltine and multivoltine and F1 varieties. After fertilization the female moths lay eggs on a piece of clean cloth or on a specially designed card board divided into several squares enclosed by cylindrical cellular (Fig. 2a). The eggs are then collected and washed in 5% bleaching to remove the gummy substance and thereafter treated with 2% formaline solution for sterilization so that the eggs may free from infectious diseases. But for protozoan infections the female moths are examined microscopically. DFL are handed over to the farmers who grow the host plant in their own farm land and use the leaves for rearing. Then the average number of DFL reared was counted and the quantity of cocoon produced (kg) in one acre farm land was also estimated (Radjabi, 2011).

### 2.9 Economic traits

#### 2.9.1 Market and selling price of cocoon

Market rate of per kg cocoons of three different varieties of silk worm was estimated. Similarly the selling price was calculated with the amount of cocoon of three different races of *B. mori*produced in per acre farm land.

Selling price of cocoon = Amount of cocoon produced in one acre farm x Rate of per kg cocoon

#### 2.9.2 Net profit of three races of B. mori

Net profit was calculated with the selling price of cocoon and deducted the cost invest on per acre of leaf producing farm land including agricultural operations *viz*. and tilling, rearingpruning, removal of dry stems and weeds and also the cost of chemical fertilizers (urea, phosphate, potash), irrigation cost, cost of seed or eggs for 500 DFL of respective variety, cost of lime needed for sterilization of the rearing room and some miscellaneous expenses.

### 2.10 Statistical analysis

By using the statistical platform National Agricultural Research System (NARS), Indian Agricultural Statistics Research Institute (IASRI), New Delhi, quantified data was analyzed. The mean difference comparison between the varieties was analyzed by analysis of variance (ANOVA) and subsequently by Tukey’s HSD(Honestly significant difference) at 5%. The heatmap was generated by Bray-Curtis distance-based redundancy analysis (dbRDA) using the Vegan package in R software version 3.5.0.

## 3. Result

### 3.1 Weight of 10 mature larvae

The average weight of 10fifth instar mature larvae of three races of *B. mori* was ranges from 28-36 gm (Fig. 3).Significantly (P<0.05) the highest weight of mature fifth instar larva was observed in bivoltine race (36 gm) and the least was recorded in multivoltine silk worm (28 gm).

**Fig. 3.**
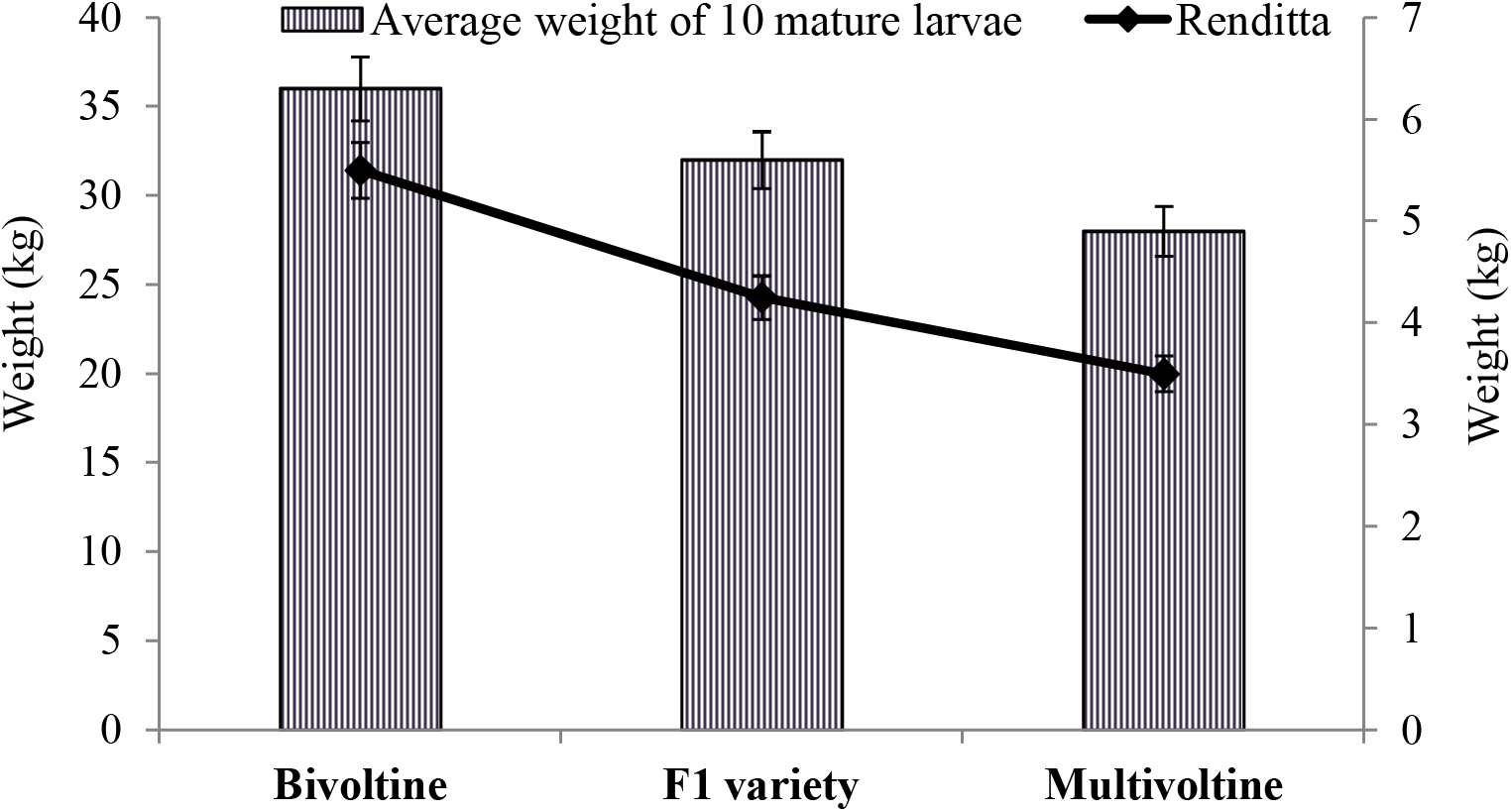
The graph represented the average weight of ten mature larvae and renditta of bivoltine, F1 and multivoltine mulberry silk worm of *B mori*.

### 3.2 Quantity of leaves required for larvae rearing

The requirement of mulberry leaves for the rearing of 100 DFL of different varieties of silk worm is shown in Fig. 4. The food consumption of bivoltine silkworm was maximum (1000 kg) and multivoltine worm consumed the least amount (600 kg).

**Fig. 4.**
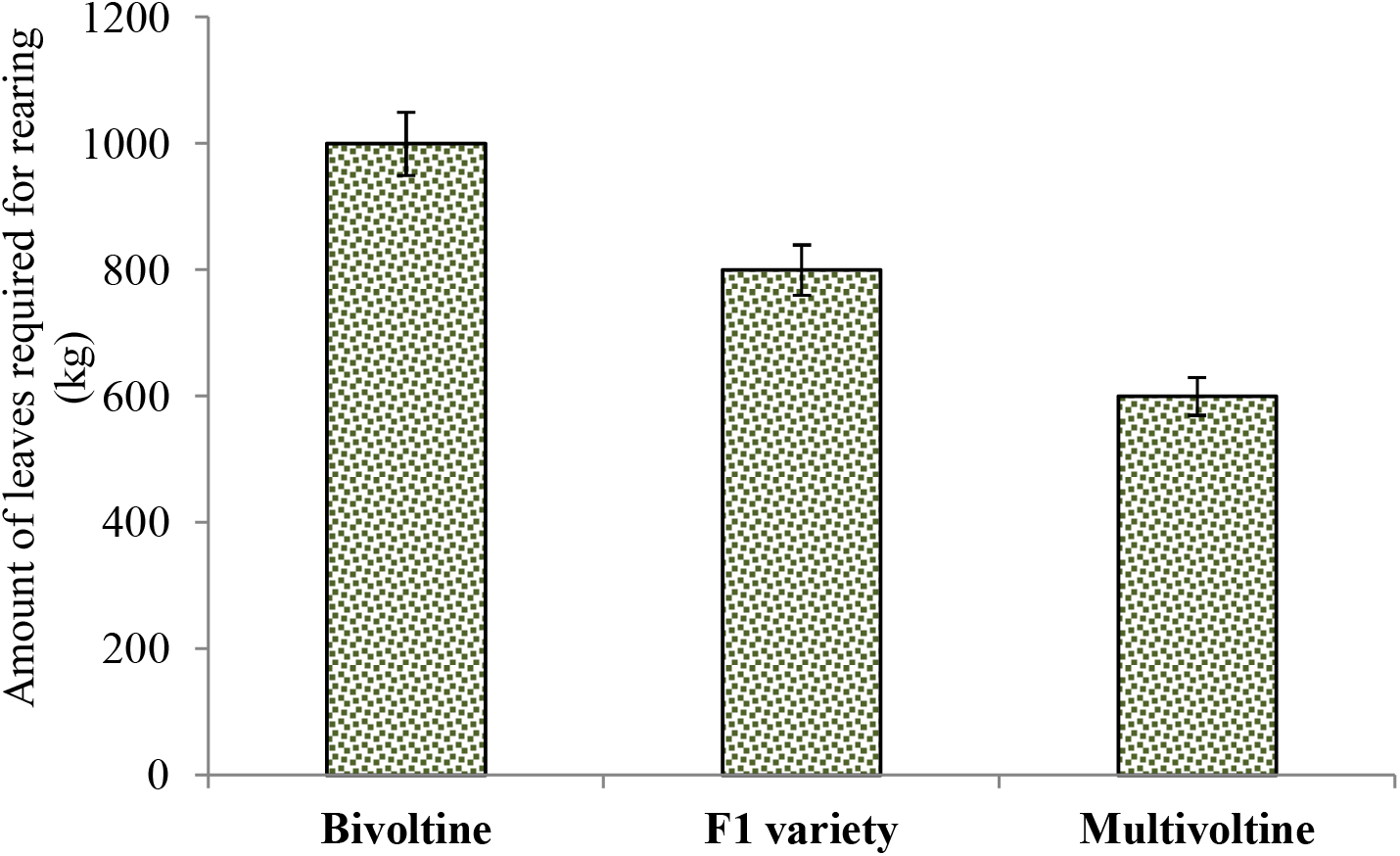
The bar graph represented the amount of leaves required for rearing the bivoltine, F1 and multivoltine mulberry silk worm of *B mori*.

### 3.3 Cocoon characters

The chhit values of different mulberry silk worm were in a range of 550-900 (unit relative) (Fig.5). The chhit value of multivoltine cocoon was maximum (900) and least was recorded in bivoltine races *B. mori* (550). Whereas, of weight of a F1 cocoon was showed moderate compared to the others.

**Fig. 5.**
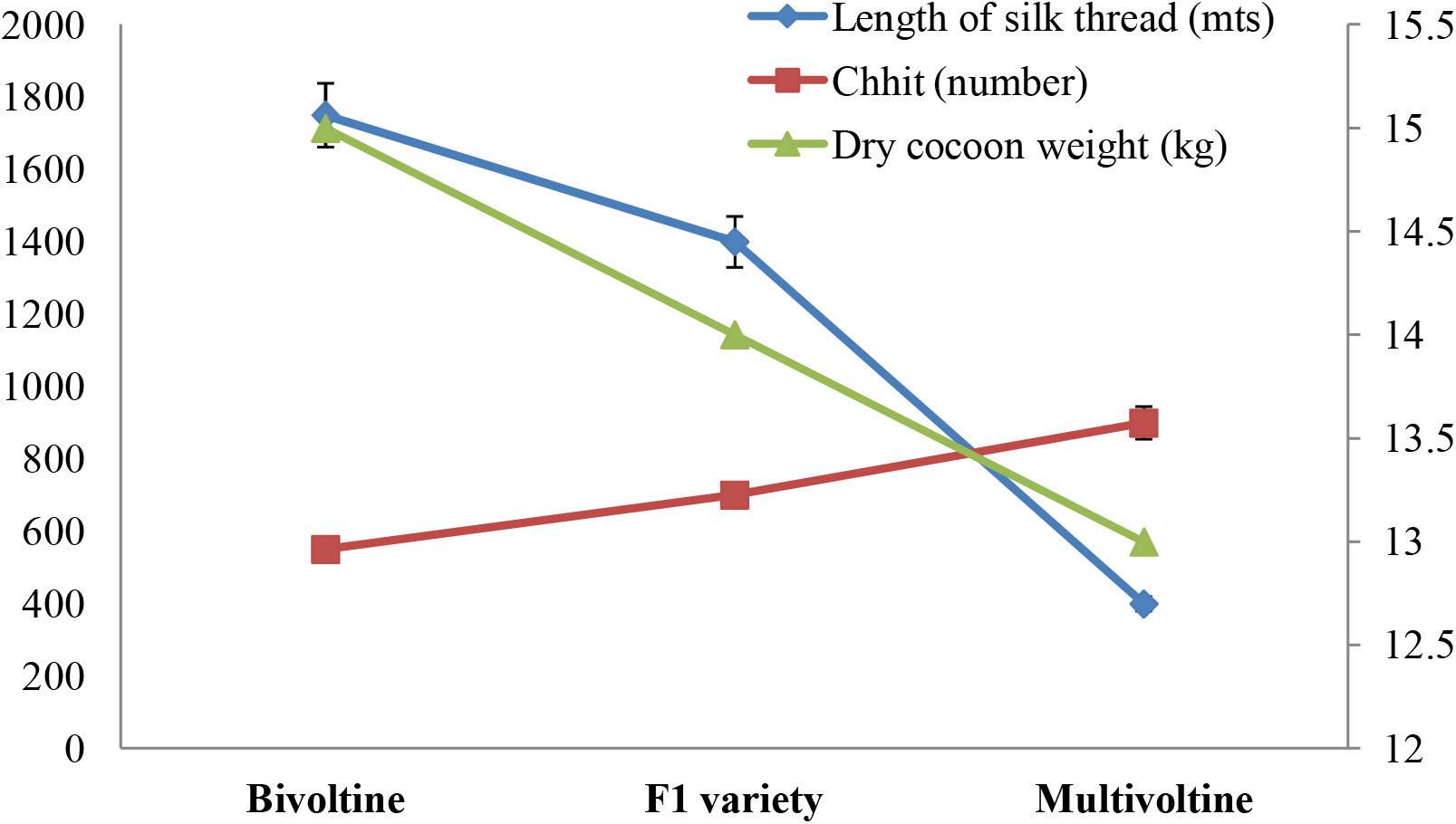
The line graph represented the length of the silk thread; chhit, number of green cocoons weigh per kg; dry cocoon weight in bivoltine, F1 and multivoltine mulberry silk worm of *B mori*.

The dry cocoon weight of 40 kg green cocoon was harvested from different variety was showed in Fig. 5. 40 kg green cocoon of different variety were dried and recorded the highest weight in bivoltine (15 kg) and the least was recorded in multivoltine races of *B. mori* (13 kg).

### 3.4 Post cocoon characters

The average length of silk thread extracted from a single cocoon of multivoltine silk worm was recorded the least length *i.e*. 400 m and significantly (P<0.05) the longest silk thread was obtained from the bivoltine cocoon *i.e*. 1750 m. whereas, silk thread length from a cocoon of a F1 variety wasmoderate (Fig. 5).

The average amount of silk thread produced from 40 Kg. of different varieties of dry cocoonswas ranges from 3.5-5.5 kg. The silk thread extracted from a constant amount of dry cocoon was recorded maximum in bivoltine cocoon (5.5 kg), and minimum was in multivoltine cocoon (3.5 kg).

### 3.5 DFL reared and cocoon produced per acre farm land

In one acre of farm land the maximum number of DFL was reared belonged to multivoltine races of *B. morii.e500* and the minimum number DFL reared belonged to bivoltine variety (200) (Fig. 6). Similar trend was observed in cocoon production in one acre farm (Fig. 6). The highest amount of cocoon produced from multivoltine (200 kg) where, the lowest amount was observed in bivoltine variety (120 kg) (Fig. 6).

**Fig. 6.**
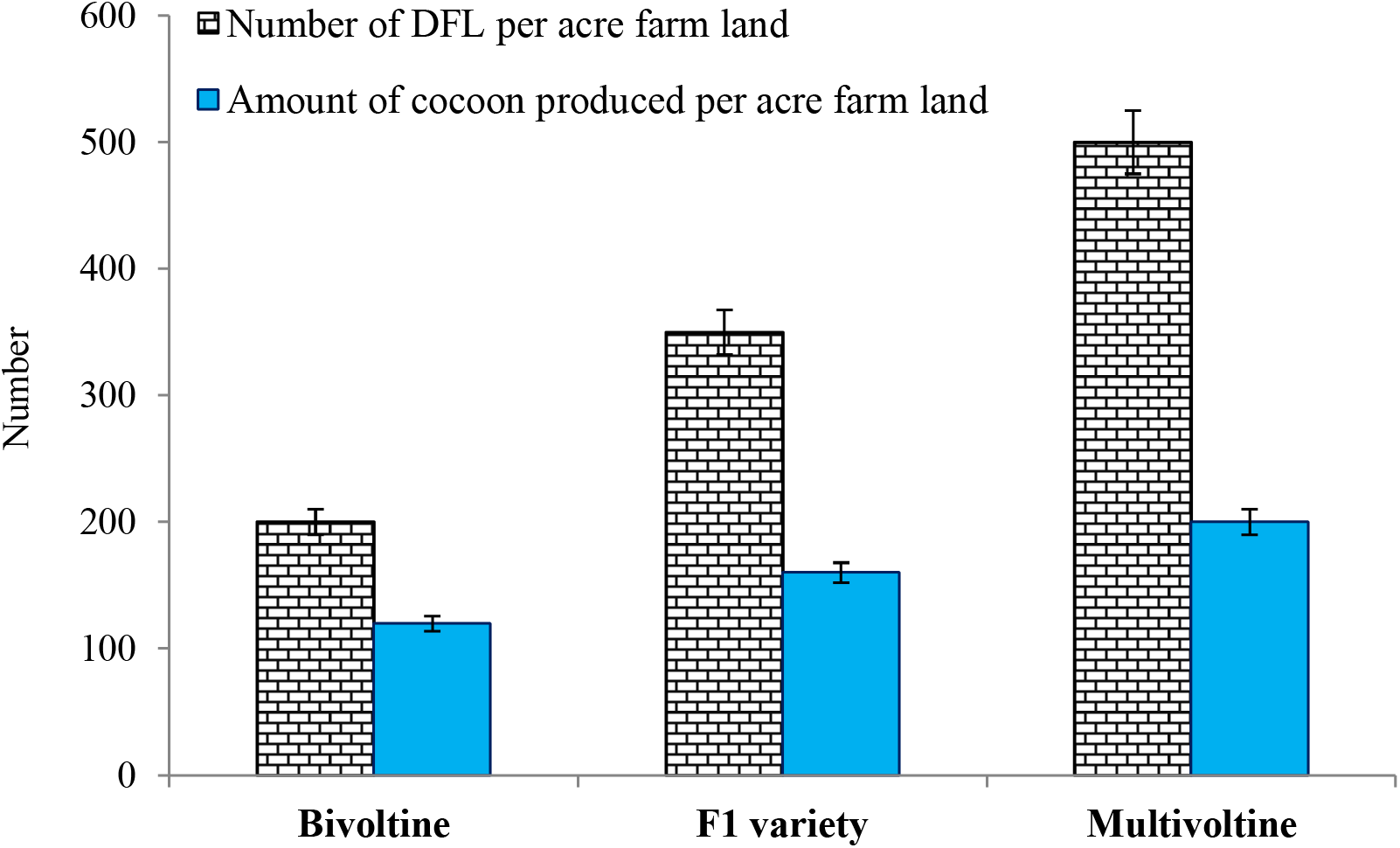
The bar graph represented the number of disease free larvae (DFL) rearing on per acre farm land and amount of cocoon produced per acre farm land in bivoltine, F1 and multivoltine mulberry silk worm of *B mori*.

### 3.6 Economy

The average market price of one kg. of silk was extracted from different varieties of silk worm showed in Fig. 7. We found that the market rate of bivoltine cocoon was the highest (500 in rupees) and that of the muiltivoltine cocoon was the least (Rs. 270/-). Whereas the selling price was calculated as per the cocoon produced in one acre farm land and the maximum selling price was observed in bivoltine followed by F1 and multivltine variety i.e. Rs. 60,000/-, Rs. 56,000/- and Rs. 54,000/-, respectively. The net profit ranges from Rs. 34,260 - 40,620/-, the highest net profit coming from bivoltine races of *B. mori* than the multivoltine and F1 varieties.

**Fig. 7.**
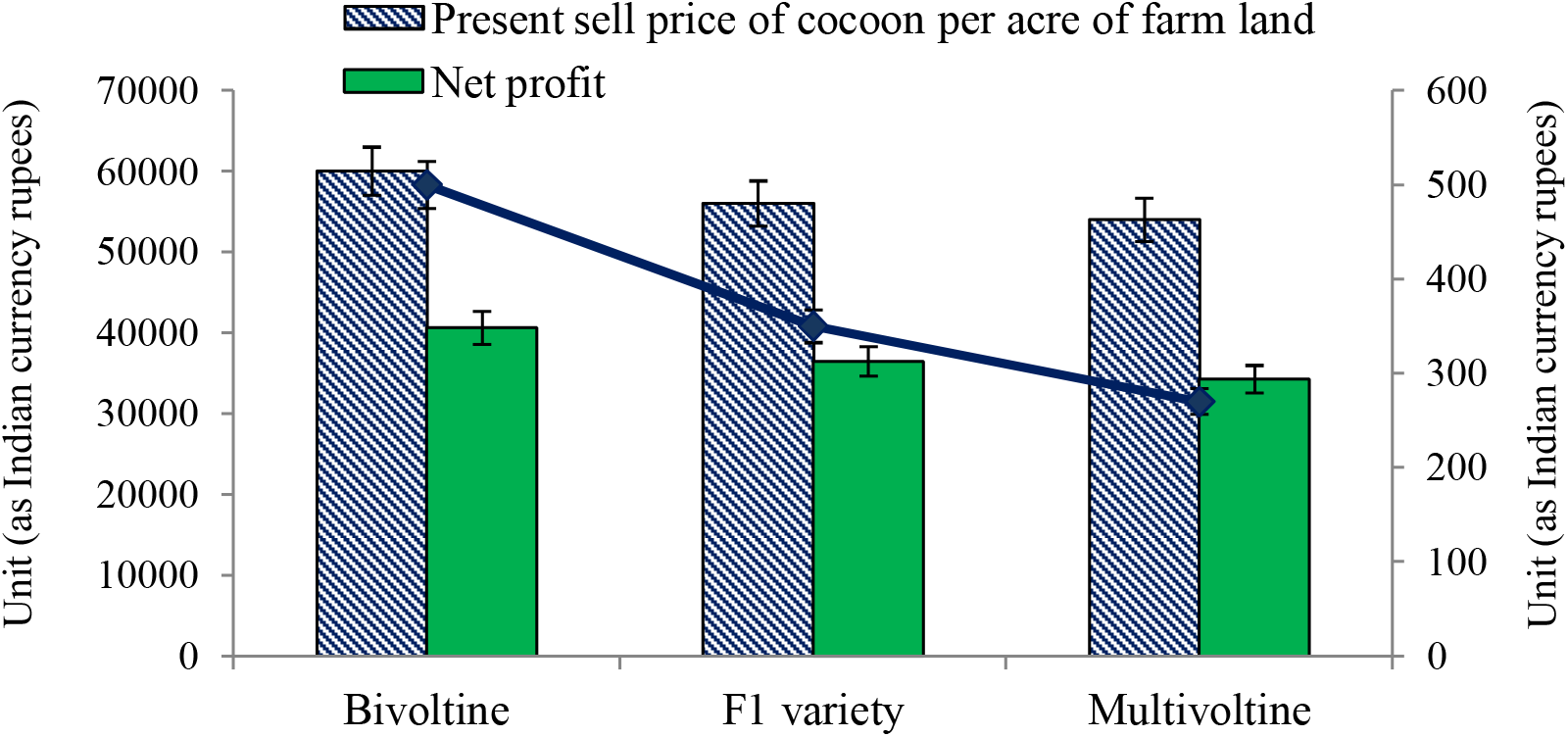
The graph represented the economic traits of bivoltine, F1 and multivoltine races of *B. mori* in terms of present selling price of respective cocoons, present market rate of cocoons and net profit per acre farm land.

### 3.7 Heatmap analysis

Heatmap is a 2D graphical color image representation of data which makes use of a predefined color scheme, and different colors display different values and variations in a data matrix (Fig. 8). The values between 1 and 2 showed the highest values with respect to the presence of color, whereas −0.5 to −1 and its relevant color showed the lower values.Here, heatmap revealed that the maximum taken parameters are positively correlated with the race of bivoltine mulberry silk worm and the most responsible variables were namely length of silk thread (LT), renditta, dry cocoon weight (DCW), average weight of 10 mature larvae (AWM), present market price (MP), leaves required for rearing, net profit and selling price of cocoon per acre farm land (SPC). In Fig. 8 the bivoltine races of *B. mori* was separately clustered from the multivoltine and F1 variety, where this two clustered unite.

**Fig. 8.**
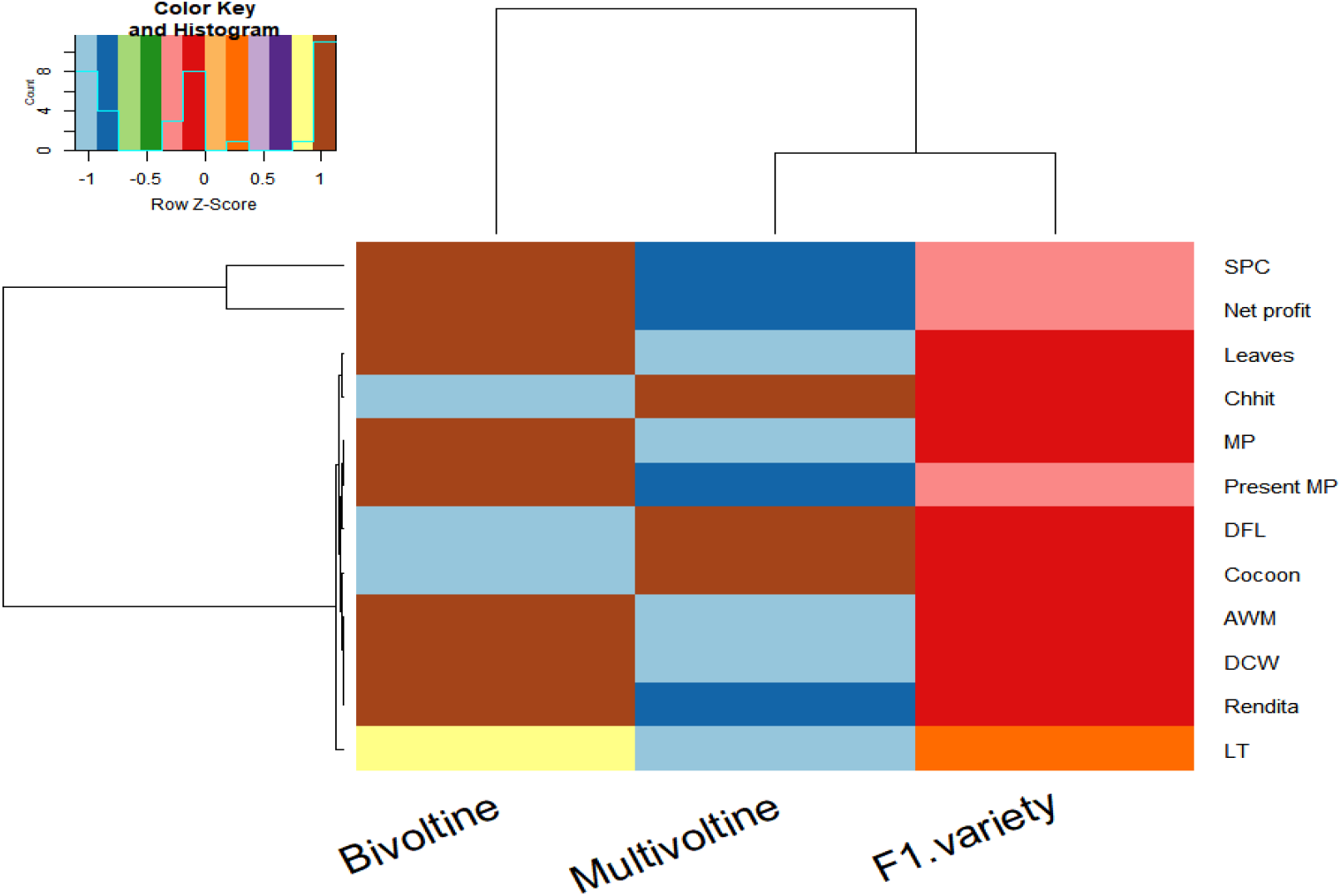
The heatmap visualized different measured variables with bivoltine, F1 and multivoltine races of *B. mori*. The color code from Z score 0-1 depicted higher values whereas, color code from Z score 0 to −1 showed lower values. Variables (LT-length of the silk thread, Renditta, DCW-dry cocoon weight, AWM-average weight of ten mature larvae, cocoon-amount of cocoon produced in one acre farm land, DFL-number of disease free larvae per acre farm land, MP-present market price of cocoon, Chhit, mulberry leaves required for larvae rearing, net profit and SPC-present sell price of cocoon per acre farm land) and varieties are grouped according to their similarity cluster in profile.

## 4. Discussion

The present investigation was made on comparative study of bivoltine, multivoltine and F1 variety of mulberry silk worm *Bombyx mori*. Before going to the field study we studied details about the optimal environmental condition for mulberry silk culture particularly during rearing period (Table 1), which is an essential to derive better financial gain. Besides, optimal temperature and humidityskilled manpower is also vital in handling the larvae during the entire rearing period.

Mulberry (*Morus* L.) is essential for the sericulture industry as the primary feed for the silkworm*B. mori*. The foliage of the host plant (*Morus indica*, primary host plant) should be soft. The larvae specially the newly hatched ones choose the young and tender leaves and thereby their growth is very likely to be optimum.Since the mulberry silk worms are totally domesticated, they have to be reared on indoor basis. So, there is no possibility to be swallowed by the birds for which the farmers have to face huge financial loss. Mating was occurred in captivity located in the farm for bivoltine, multivoltine and F1 varieties. After fertilization the female moths lay eggs on a piece of clean cloth or on a specially designed card board divided into several squares enclosed by cylindrical cellular. The eggs are then collected, washed and sterilized for free from infectious diseases. But for protozoan infections the female moths are examined microscopically. DFL are handed over to the farmers who grow the host plant in their own farm land and use the leaves for rearing.

The present study showed that the tenmature larvalweights of bivoltine races of *B*. *mori* was higherto that of the multivoltine and F1 variety. Highlighting the importance of food intake, Horie *et al*. (1978) reported that requirement of ingestion and digestion of food is 4.2 mg and 1.8 mg, respectivelyfor the production of 1 g larval dry weight. The intake of food during total larval life gets reflected by the weight of 10 mature larvae (Ito, 1978). Maximum amount of leaves were required for rearing of bivoltine variety followed by F1 and multivoltine race. In the present study, findings were similar as reported by Rao *et al*. (1998).Silkworm fecundity is also affected by feeding quality and mulberry varieties (Table 2). Number of laid eggs of bivoltine variety much healthier than the multivoltine and F1 varitywhich achieved due to fresh leaves consumption.Previously researchers also reported the same on the aspect on the effect of mulberry varieties on fecundity, silk glands and cocoon characteristics (Krishnaswami et al., 1970; Krishnaswami et al., 1971; Legay, 1958)

**Table 2.** Estimation of fecundity in multivoltine silk worm at optimal temperature 22-28°C and relative humidity of the atmosphere 60-85%. Fecundity= Number of eggs lay by each female moth immediately after mating. Here fecundity calculated by total number of eggs laid on a specialized card board demarcated with twenty squares for 20 female moths, which surrounded by a cylindrical cellule (Fecundity= Total number of eggs in twenty cellule (9679) / Number of female months laid eggs (20).

| Number of eggs | 1<sup>st</sup> cellule | 2<sup>nd</sup> cellule | 3<sup>rd</sup> cellule | 4<sup>th</sup> cellule | 5<sup>th</sup> cellule | Total | Avg. fecundity |
| --- | --- | --- | --- | --- | --- | --- | --- |
| Row 1 | 490±10 | 380±8 | 450±10 | 310±5 | 650±10 | 2280 | 484 |
| Row 2 | 710±5 | 240±10 | 315±8 | 414±10 | 680±10 | 2359 |  |
| Row 3 | 430±10 | 730±10 | 460±10 | 490±10 | 210±8 | 2320 |  |
| Row 4 | 340±10 | 730±10 | 355±8 | 720±10 | 475±10 | 2620 |  |

The cocoon characters are important components, which determine the overall performance of the silkworm and also important for subsequent silk fiber quality (Kumar et al., 2012; Malik et al., 2006; Joge et al., 2003). The result revealed that the bivoltine races showed better performance statistically (P<0.05) in terms of cocoon production compared to the multivoltine and F1 variety, which is corroborated with the recent reports of Sharma and Bali, (2019). They showed the bivoltine cocoons were much superior in comparison to multivoltine with higher silk content, longer filament, higher neatness, cleanness, low boil off loss ratio, higher tensile strength and less variation in evenness(Sharma and Bali, 2019; Dandin et al., 2003).Similar trends of result found in average length of silk thread from a single cocoon of bivoltine races, which was higher by 30% and 77.14% from F1 and multivoltine variety, respectively. The probable reason might be dietary changes in the silkworm, which directly or indirectly influence on the cocoon characters and on post cocoon characters such as quantity of the silk filament and denier (Malik and Reddy, 2007).Various factors such as genetic background of silkworm, food quality, environment and etc., influence the economic traits of the silkworm (Lokesh and Anantha, 2011; Raina, 2001).

In the present study we found that the market rate of bivoltine cocoon was the highestthat of the multivoltine cocoon. Whereas the selling price was calculated as per the cocoon produced in one acre farm land and the maximum selling price was observed in bivoltine followed by F1 and multivoltine variety respectively. The net profit for bivoltine races of *B. mori* was significantly increased by 15.65% and 10.21% in multivoltine and F1 varieties, respectively. From these accounts of expenditure, it was found that other than the cost of DFL, remaining expenses were constant for the procurement of a single crop and this variation for DFL was nominal. But in case of multivoltine silk worm it yields five crops in a year and simultaneously the expenditure on food consumption will be increased proportionally. Since there were no diapauses in this variety, the adult worm immerges from the cocoon almost immediately after cocoon formation completed. Thereafter they undergo coupling, after which the female moth starts to lay eggs. Mulberry leaves required for the rearing of those eggs cannot be available from the same farm land which has already been exhausted during the rearing of first crop. So, another farm land of same area is necessary and accordingly the expenditure will be enhanced.Whereas in case of bivoltine silk worm, per annum two crops are procured and there is enough time gap in between two crops. So the mulberry leaves for feeding the larvae can be easily available from the same one acre field, if irrigation, manuring, pruning and other maintenance job can be done properly. Therefore, the overall expenditure will be much less in comparison with the multivoltine silk worm.Moreover, the heatmap analysis revealed that the bivoltine races of *B. mori* was separately clustered based on the positively correlation of measured variables mostly length of silk thread, renditta, dry cocoon weight, average weight of 10 mature larvae, present market price, leaves required for rearing, net profit and selling price of cocoon per acre farm land; whereas, the multivoltine and F1 variety clustered in a separate group.Therefore, from the above investigations on the aspect of correlationfound that the bivoltine variety is much superior and also better from the angle of economical as well as biological perspective.

## 5. Conclusion

In the present study, results showed the comparison of biological and economic traits of bivoltine, multivoltine and F1 races of *B. mori* and concluded that silk produced from the cocoons of bivoltine variety is much more superior in both the quantity and quality wise than other two races. For multivoltine race, generally five crops can be obtained in a year and there is almost no time gap to prepare the food plants for the rearing the next crop. So, if any farmer like to rear all the five generations in a year, he has to provide extra farm land at two acre which enhances the total investment. But in case of bivoltine race, where two life cycles complete in a year, thereby, there is enough time gap in between two consecutive voltines, within which the same land can yield the mulberry leaves required for feeding the larvae, if it is maintained properly, i.e., irrigation, manuring, pruning and other maintenance tasks. Besides, the silk extracted from the cocoons of the bivoltine variety is superior both in quantity and quality to that of the multivoltine type. So, the families who are involved in silk worm rearing should choose the bivoltine silk worm for better monetary gain at the cost of less labor and other maintenance cost. The silk industry in West Bengal as well as in India is in sick and it provides subsidiary income to the families involving in silk worm rearing specially the women folk. So, attention should be given to educate and motivate them specially the poor and tribal people living in arid zone of the country by community awareness. This comparative study suggests that sericulture particularly with bivoltine races of *B. mori* is capable of more income generation than other traditional agricultural crops.

## 6. Abbreviations

DFL: Disease free larvae
ERR: Effective rate of rearing
LT: Length of silk thread
DCW: Dry cocoon weight
AWM: Average weight of 10 mature larvae
MP: Present market price
SPC: Selling price of cocoon per acre farm land

## 7. Declarations

### 7.1 Ethics approval and consent to participate

Not applicable.

### 7.2 Consent for publication

Not applicable.

### 7.3 Availability of data and materials

All the data and materials presented in the manuscript are the original work of the authors.

### 7.6 Authors’ contributions

MK and KN performed the experiment. MK has written the manuscript and analyzed the data. MK, UK, SNC, SLS, MS and KN read and approved the final manuscript.

## Acknowledgments

We thank all the erstwhile researchers associated with this research. The head of the department of Zoology, Asansol Girl’s College, West Bengal, is highly acknowledged for providing necessary facilities to carry out this research. We also acknowledge ICAR-National Rice Research Institute, Cuttack, Odisha, for analytical support.

## References

Adolkar, V.V., Raina, S.K., Kimbu, D.M., 2007. Evaluation of various mulberry *Morus* spp. (Moraceae) cultivars for the rearing of the bivoltine hybrid race Shaanshi BV-333 of the silkworm *Bombyx mori* (Lepidoptera: Bombycidae). International Journal of Tropical Insect Science, 27(1): 6–14.

Anonymous, 2016. Seridoc dairy 2016, Govt. of West Bengal.

Brahma, D., Swargiary, A., Dutta, B., 2015. A comparative study on morphology and rearing performance of *Samia ricini* and *Samia canningi* crossbreed with reference to different food plants. Journal of Entomology and Zoology Studies, 3(5): 12–19.

CSB. 2018. Note on the performance of Indian silk industry & functioning of Central Silk Board, 28–29.

Dandin, S.B., Basavaraja, H.K., Suresh, Kumar, N., 2003. Factors for success of Indian bivoltine sericulture. Indian Silk. 41(9):5–8.

Doddaswamy, M.S., Subramanya, G., Talebi, E., 2009. Studies on some economic traits and biological characters of regular and reciprocal cross between a multivoltine and bivoltine race of the silkworm *Bombyx mori*. Journal of Entomology and Nematology. (4):50–55.

Gangwar, S.K., 2010. Impact of varietal feeding of eight Mulberry varieties on *Bombyx mori* L., Agriculture and Biology Journal of North America, 1(3): 350–354.

Horie, Y., Inokuchi, T., Watanabe, K., 1978. Quantitative studies of food utilization by the silkworm *Bombyx mori* L. through the life cycle II. Economy of nitrogen and amino acids. Bulletin of Sericultural Experimental Station. 27(2):531–578.

Ito, T., 1978. Silkworm Nutrition; in the Silkworm an Important Labratoary Tool. Tazima, Y. (Ed), 121–157, Kodansha Ltd, Tokyo.

Joge, P.G., Pallavi, S.N., Begum, N.A., Mahalingappa, K.C., Mallikarjuna, Mishra, R.K., Gupta VP et al. 2003. Evaluation of double hybrids of silkworm *Bombyx mori* L. in the field. In: Advances in Tropical Sericulture. Dandin, S.B., Mishra, R.K., Gupta, V.P., Reddy, Y.S., (eds.), 102–104, NASSI, Bangalore.

Jolly, M.S., Ed. 1987. Appropriate sericulture techniques ICTRETS, Mysore, India. 75.

Kamili, A.S., Malik, G.N., Trag, A.R., Kukiloo, F.A., Sofi, A.M., 2000. Development of new bivoltine silkworm (*Bombyx mori* L.) genotypes with higher commercial characters. Journal of Research, SKUAST. 2:66–69.

Koidzumi G. 1917. Taxonomy and phytogeography of the genus *Morus*. Bull. Seric. Exp. Station, Tokyo (Japan) 3: 1–62.

Krishnaswami, S., Noamani, K.R., Asan, M., 1970. Studies on the quality of mulberry leaves and silkworm cocoon crop production, Part I, Quality difference due to varieties. Indian Journal of Sericulture, 9(1): 87–93.

Krishnaswami, S., Kumarraj, S., Vijayaragavan, K., Kasiviswanathan, K., 1971. Silkworm feeding trails for evaluating mulberry leaves as influenced by variety, spacing and nitrogen fertilization. Ibid, 5: 13–17.

Kumar, S.N., Murthy, D.P.P., Moorthy, S.M., 2011. Analysis of heterosis over environments in Silkworm (*Bombyx mori* L.). ARPN Journal of Agricultural and Biological Science. 6(3):39–47.

Kumar, S.N., Prakash, Murthy, D.P., Moorthy, S.M., 2012. Analysis of heterosis over environments in silkworm (*Bombyx mori* L.). ARPN Journal of Agricultural and Biological Science. 6(1):3–10.

Lakshmi, H., Chandrashekharaiah, Babu, M.R., Raju, P.J., Saha, A.K., Bajpai, A.K., 2011. HTO5 x HTP5: The new bivoltine silkworm *Bombyx mori* L. hybrid with thermotolerence for tropical areas. International Journal of Plant, Animal and Environmental Sciences. 1(2):88–104.

Lee, Y.W., 1999. Silk reeling and testing manual. FAO Agricultural Services Bulletin No. 136. Rome, Italy.

Legay, J.M., 1958. Recent advances in silkworm nutrition. Annual Review of Entomology, 3: 7586.

Linneaus, C., 1753. Species Plantarum, Stockholm, pp. 986.

Lokesh, G., Anantha, Narayana, S.R., 2011. Changes in the Protein profile of silkworm *Bombyx mori*. L (Lepidoptera: Bombycidae) in response to the chemical mutagen. International Journal of science nature. 2(3):559–563.

Malik, F.A., Reddy, Y.S., 2007. Role of mulberry nutrition on manifestation of post cocoon characters of selected races of the silkworm *Bombyx mori* L. Serocologia. 47:63–76.

Malik, G.N., Massoodi, M.A., Kamili, A.S., Sofi, A.M., 2006. Studies on heterosis in some bivoltine silkworm (*Bombyx mori* L.) crosses. Journal of Sericulture. 6(1-2):47–49.

Mano, Y., Kumar, S.N., Basavaraja, H.K., Reddy, N.M., Datta, R.K., 1992. An index for multiple trait selection for silk yield improvement in *Bombyx mori* L. National Conference on Mulberry Sericultural Research, Central Sericultural Research and Training Institute, Mysore, India. 116.

Masrat, S., Tripathi, A.K., 2017. Sericulture Industry in India A comparative study of Jammu and Kashmir and Madhya Pradesh, International Journal of Business and Administration Research Review. 1(18):81.

Radjabi, R., 2011. Nutritive Quality Assessment of Two Mulberry Varieties including Kokosa and Local on Some Biological Parameters and Economical Characters of Silkworm *Bombyx mori* L. Advances in Environmental Biology, 5(7): 1877–1881.

Raina, S.K., 2000. The Economics of sericulture and apiculture modules for income generation in Africa. IBRA, UK. 86.

Rao, G.S., 1998. Evaluation and authorization of parent races and their hybrids – Modalities. In: Silkworm breeding, G. Sreerama Reddy (Ed.). Proceedings of the National Workshop March 1819. Oxford and IBH publishing Co. Pvt. Ltd., New Delhi, India, 227.

Roopa, H., Murthy, C., 2015. Trends in arrivals and prices of cocoons in Shirahatti market at Haveri district. International Journal of Commerce and Business Management, 8(1): 131–134.

Sharma, K., Bali, K., 2019. Evaluation of indigenous and introduced bivoltine silkworm hybrids. Journal of Pharmacognosy and Phytochemistry, 8(4): 1459–1464.

Talebi, E., Subramanya, G., 2009. Genetic distance and heterosis through Evaluation Index in silkworm *Bombyx mori* L. World Applied Sciences. 7(9):1131–1137.

Tsukada, M., Islam, S., Arai, T., Bosch, A., Fred, G., 2005. Microwave irradiation technique to enhance protein fiber properties. Autex Research Journal. 5(1): 40–8.

Ullal, S.R., Narasimhanna, M.N., 1987. Handbook of practical sericulture. 3rd ed. Central Silk Board, Bangalore, India. 166.

